# The polyanionic drug suramin neutralizes histones and prevents endotheliopathy

**DOI:** 10.1101/2021.12.09.469611

**Authors:** Nuria Villalba, Adrian M. Sackheim, Michael A. Lawson, Laurel Haines, Yen-Lin Chen, Swapnil K. Sonkusare, Yong-Tao Ma, Jianing Li, Dev Majumdar, Beth A. Bouchard, Jonathan E. Boyson, Matthew E. Poynter, Mark T. Nelson, Kalev Freeman

## Abstract

Drugs are needed to protect against the neutrophil-derived histones responsible for endothelial injury in acute inflammatory conditions such as trauma and sepsis. Heparin and other polyanions can neutralize histones but may cause secondary, deleterious effects such as excessive bleeding. Here, we demonstrate that suramin—a widely available polyanionic drug—completely neutralizes the toxic effects of histones. The sulfate groups on suramin form stable electrostatic interactions with hydrogen bonds in the histone octamer with a dissociation constant of 250 nM. In cultured endothelial cells (Ea.Hy926), histone-induced thrombin generation was significantly decreased by suramin. In isolated murine blood vessels, suramin abolished aberrant endothelial cell calcium signals and rescued impaired endothelial-dependent vasodilation caused by histones. Suramin significantly decreased pulmonary endothelial cell ICAM-1 expression and neutrophil recruitment caused by infusion of sub-lethal doses of histones *in vivo*. Suramin also prevented lung edema, intra-alveolar hemorrhage and mortality in mice receiving a lethal dose of histones. Protection of vascular endothelial function from histone-induced damage is a novel mechanism of action for suramin with therapeutic implications for conditions characterized by elevated histone levels.

**Significance Statement:** Pathologic levels of circulating histones cause acute endotheliopathy, characterized by widespread disruption of critical endothelial functions and thromboinflammation. We discovered that suramin binds histones and prevents histone-induced endothelial dysfunction, thrombin generation, lung injury, and death. Histone binding is a novel mechanism of action for suramin, considered among the safest and most effective drugs by the World Health Organization. These results support the use of suramin for protection of blood vessels in conditions exacerbated by circulating histones including trauma and sepsis.

## Introduction

Acute endotheliopathy is a clinical syndrome resulting from extensive tissue injury in trauma and sepsis, including that attributable to SARS-CoV-2 infection. Endotheliopathy is characterized by widespread disruption of endothelial-dependent vasodilatory function, barrier integrity, and hemostasis which all contributes to thromboinflammation, organ failure, and mortality (1, 2). Extracellular histones are major mediators of endotheliopathy, as shown by the efficacy of anti-histone antibodies in preventing systemic inflammation and mortality in animal models of sepsis and endotoxemia through lipopolysaccharide (LPS) infusion (3, 4). Histones enter the circulation when released by cellular apoptosis or necrosis (5–7), and in innate immunity, when activation of neutrophils leads to the release of chromatin in the form of neutrophil extracellular traps (NETs). These NETs contain granular enzymes and peptides which aid in clearing bacteria, and nuclear proteins, predominately histones (3, 4). Nucleosomes induce cytokine production at low concentrations, but high concentrations kill cells (8). Evidence of NET-induced endothelial damage has been reported in COVID-19 (9), atherosclerosis (10, 11), ischemia/reperfusion (12), and venous thrombosis (13, 14). Plasma nucleases act on DNA-histone complexes circulating in the blood to degrade the nucleic acids, exposing the highly cationic histones that function as damage-associated molecular pattern proteins, activating the immune system and causing additional toxicity (15–17). Free histones are found at low levels (2-5 μg/mL) in the circulation in uninjured humans, but levels can reach 20-100 μg/mL in COVID-19 (18) and up to 250 μg/mL in the acute period following severe trauma before they are degraded over hours and days by the protease activated protein C (19). At high concentrations, histones can activate platelets and damage vascular cells, particularly pulmonary (19) and mesenteric endothelial cells (20), vascular smooth muscle cells (11) and erythrocytes (21, 22), but histones are not directly toxic to endothelial cells in the cerebral vasculature (23). Histones activate and injure endothelial cells through mechanisms including calcium overload (3, 20), pyroptosis through NLRP3 inflammasome and toll-like receptor activation (TLR) (6, 24–27), and disassembly of adherens junctions causing loss of barrier function (23, 28). Histones, like other cations can also bind directly to anionic membrane phospholipids in stoichiometric ratios (29, 30), with high concentrations leading to disruption of lipid bilayers (11, 31). Furthermore, histones deposited on the lumen of blood vessels can also attract monocytes in a surface-charge dependent fashion causing atherosclerosis (4). Elevated histone levels have been linked to widespread endothelial injury and organ damage in human patients after trauma (19, 32–38) and other conditions including ischemic stroke (39), sepsis (3), pancreatitis (40), and acute respiratory distress syndrome (ARDS) (41, 42).

The critical unmet need for therapeutics that protect the vascular endothelium from histone-mediated injury has become of immediate relevance in the context of the SARS-CoV-2 pandemic (1, 43). We previously used native blood vessels from humans and mice to show the spatio-temporal distribution of endothelial cell calcium signals after histone exposure (20). Using resistance-sized mesenteric arteries from mice expressing an endothelial cell-specific fluorescent calcium biosensor, we found that histone-induced calcium signals were blocked by the removal of extracellular calcium, but unaffected by depletion of intracellular calcium stores with cyclopiazonic acid (CPA). Because TLR receptors have been implicated in histone effects in platelets and other cells (6, 25, 44), and TLR agonists can trigger store-operated calcium (SOC) entry in endothelial cells (45), we suspected that TLR activation might explain histone-induced calcium influx. However, we and others have shown that TLR2 and TLR4 are not significantly involved in endothelial calcium entry or cytotoxicity induced by histones (19, 20, 46). We then hypothesized that histone-induced calcium signals could involve the non-specific cation channel transient receptor potential vanilloid 4 (TRPV4), a major mechanism of calcium entry in endothelial cells (47), but TRPV4 knockout did not diminish endothelial histone-induced calcium signals (20).

Purinergic signaling is a central component of inflammation and regulates responses to extracellular nucleotides. During inflammation, adenosine 5’-triphosphate (ATP) released from damaged tissues, dying cells or in response to stimuli via pannexins can activate endothelial purinergic receptors to trigger IP_3_ evoked calcium release, a known mechanism of pulmonary vascular inflammation (48–50). This led us to hypothesize that purinergic receptors might play a role in histone-induced events. In pilot studies we found that the non-specific purinergic receptor antagonist suramin caused a complete block of histone-induced calcium signals in microvascular endothelial preparations, but genetic ablation of P2X7 did not prevent these responses. The molecular structure of suramin (Fig. 1*A*) immediately suggested another potential explanation for our observations – it is a polyanion with the potential to bind cationic histones directly in solution. This led us to conduct fluorescence spectroscopy experiments which showed that suramin binds to histones in solution and calculated the dissociation constant (K_d_) of the histone-suramin complex. Molecular modeling also demonstrated the likely binding sites between histones and suramin (Fig 1).

**Figure 1.**
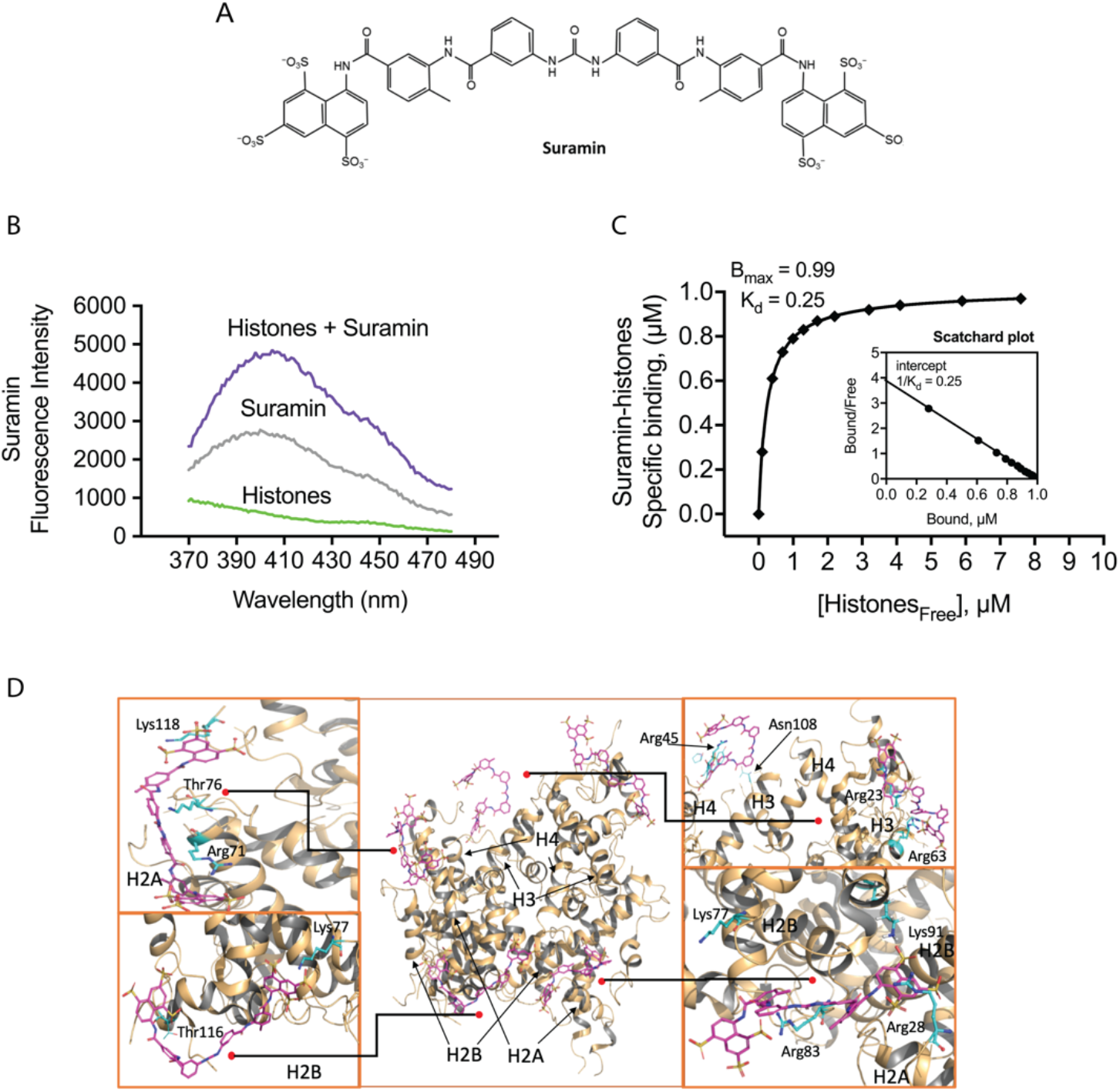
Suramin binds histones in solution. (A) The chemical structure of suramin. (B) *In vitro* fluorescent spectroscopy studies were used to biochemically establish the interaction between suramin and histones. We established the absorbance and emission spectra for histones and suramin in solution (Supplemental Fig. 1 *A*), and then measured suramin sodium salt intrinsic fluorescence using increasing concentrations of suramin to determine the saturation range of the detector (Supplemental Fig. 1 *B*). (C) Scatchard plot analysis of the binding curve demonstrates a single high-affinity binding site with a dissociation constant (K_d_) of 250 nM. (D) Molecular dynamics simulations showing interactions between suramin molecules and the histone octamer in solution. Several exposed amino acid residues including arginine, asparagine, lysine, and threonine form hydrogen bonds with the sulfate groups on suramin (Lys118, Thr76, Arg71, Lys77, Thr116, Arg45, Asn108, Arg23, Arg63, Lys91, Arg83, Arg28). These include residues on H2A, H2B, H3, and H4, which are predicted to form stable electrostatic interactions with the sulfate groups on suramin.

These unexpected observations were interesting, because suramin is not only a useful tool in pharmacology, but also a safe and widely available drug. First synthesized by Bayer in 1917 as part of a drug discovery program for trypanosomiasis (African sleeping sickness), suramin is a bis-polysulfonated naphthylurea hexaanion with activity against trypanosomes in both animal models and humans (51). Suramin has been used clinically for over 100 years as an anti-parasite and anti-cancer agent, and, importantly, is considered among the safest and most effective drugs for health care by the World Health Organization. Unlike heparan sulfate or heparin synthetic polyanions, which also bind histones, suramin dosing is infrequent (usually once per week), and is inexpensive, well-tolerated, and does not cause complications associated with anticoagulation.

Together, the published literature and our preliminary data provided a strong rationale for us to investigate suramin as a candidate drug to prevent histone-induced endotheliopathy. The objective of this study was to test the hypothesis that suramin can protect against histone-induced endothelial dysfunction. We demonstrate that histones activate human endothelial cells to promote rapid thrombin generation, a reaction that is blocked by suramin. In pressurized murine vascular preparations, we directly tested the efficacy of suramin for preventing histone-induced aberrant endothelial calcium signaling and vasodilatory dysfunction. In animal models of trauma, it is the lung tissue and not kidney or liver, which is the primary target for circulating damage-associated molecular pattern proteins (19). In a histone infusion model, we found that suramin prevented histone-induced lung injury, endothelial cell activation, adhesion molecule expression, and pulmonary barrier disruption. Importantly, we also show that suramin completely protects against a lethal dose of histones *in vivo*. Thus, histone binding is a novel mechanism of action for suramin, providing a rationale for its use to protect the endothelium in conditions exacerbated by circulating histones, such as trauma and sepsis. Together, these experiments provide new insights into the deleterious effects of histones on endothelial function, thrombin generation, and lung damage, and provide support for the use of suramin as a strategy to protect against histone-induced endotheliopathy.

## Results

### Suramin binds histones in solution

Based on its molecular structure (Fig. 1 *A*), we hypothesized that suramin, a highly charged polysulfonated napthylurea, would bind avidly to cationic histone complexes. When NETs or nucleosomes enter the bloodstream, they are exposed to endogenous nucleases that rapidly digest DNA, leaving free histone proteins (15). Therefore, we focused on testing the interactions between suramin and histones. First, fluorescent spectroscopy studies were used to biochemically establish the dissociation constant (K_d_) and number of high-affinity binding sites for interactions between the two molecules. We established the absorbance and emission spectra for histones and suramin in solution, and then measured suramin sodium salt intrinsic fluorescence using an excess of suramin in the presence of increasing concentrations of histones (Fig 1 *B;* Supplemental Fig. 1). The resulting interactions are represented using a binding curve (Fig. 1 *C*). Scatchard analysis of the binding curve demonstrates a single high-affinity binding site with a dissociation constant (K_d_) of 250 nM (Fig. 1 *C*). These results confirm that suramin readily binds histones in solution. We then used all-atom molecular dynamics (MD) simulations to determine likely interactions between suramin molecules and the histone octamer in solution (Fig. 1 *D* and Supplemental Video 1). Suramin quickly formed electrostatic contacts between its SO3- and arginines (Arg) on the protein surface such as Arg53 and Arg69 of H3, Arg23 and Arg45 in H4, Arg17 in H2A and Arg30 in H2B. Hydrogen bonding between suramin and several threonines (Thr) was also observed, such as Thr80 of H3, Thr16 and Thr76 in H2A, and Thr116 in H2B. These interactions remained stable toward the end of our simulations and enabled steady binding for five of the suramin molecules to histones.

### Histones induce rapid thrombin generation on human endothelial cells that is blocked by suramin

Thrombin is the ultimate protease in the clotting cascade, catalyzing fibrin formation. The formation of thrombin following cleavage of prothrombin is the rate limiting to the coagulation process (52). Phosphatidylserine-dependent prothrombin activation on the endothelial surface leads to the formation of microthrombi, shedding of extracellular vesicles and glycocalyx, neutrophil migration, and efflux of water into damaged interstitial tissues (10, 52). Extracellular vesicles, enriched with histones, have procoagulant membranes and carry microRNAs into the bloodstream (53). To test whether histones promote thrombin generation, we used calibrated automated thrombograms in recalcified pooled healthy human plasma, in the absence of exogenous tissue factor and phospholipid membrane, to measure the ability of cultured human endothelial cells (Ea.hy926) to support thrombin production. Under these conditions, thrombin formation occurred slowly (lag time >15 minutes), but when histones were applied, thrombin generation was accelerated, with a lag time of <5 minutes (Fig. 2 *A* and *B*). Histone treatment also had prothrombotic effects on other measures of thrombin generation including peak thrombin, endogenous thrombin potential, time to peak thrombin, and the rate of thrombin generation (Fig. 2 *C-F*). In the presence of suramin, measures of histone-induced thrombin generation were significantly ameliorated to levels observed in untreated cells.

**Figure 2.**
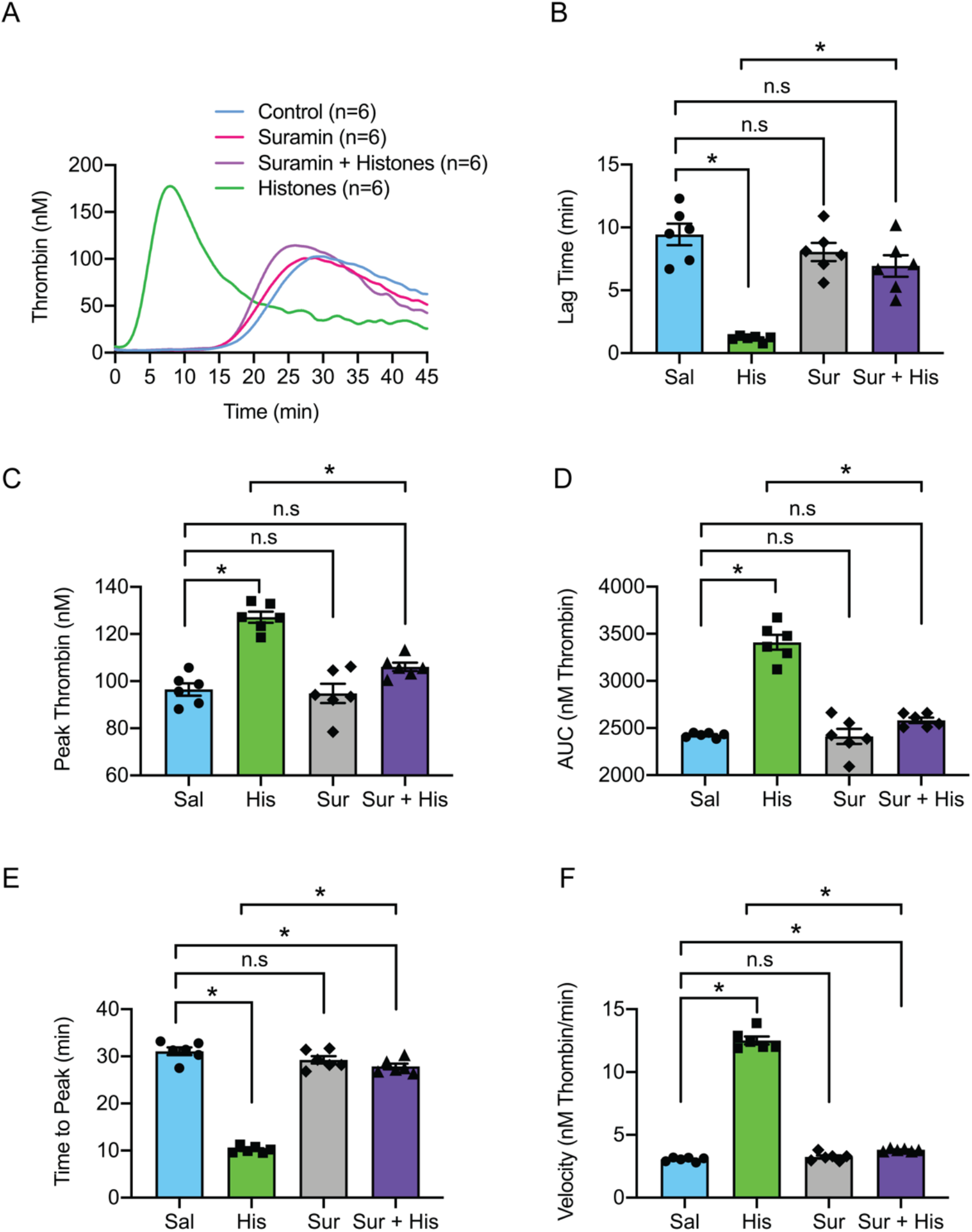
Histones drive rapid thrombin generation on human endothelial cells which is blocked by suramin. (A) Calibrated automated thrombogram (CAT) tracings of thrombin generation (nM) vs time (min) by cultured human endothelial cells (Ea.hy926) in re-calcified, pooled, healthy human plasma. Histones (50 μg/mL), suramin (50 μM) or a combination of both were exogenously added to the cell culture and plasma samples as needed. (B) Summary data for lag time (min) in control (9 ± 0.8 min; n=6), suramin (8 ± 0.7 min; n=6), suramin and histones (67 ± 0.9 min; n=6) and histones (1 ± 0.1 min; n=6) samples. (C) Summary data for peak thrombin (nM) in control (96 nM; n=6), suramin (94 ± 4 nM; n=6), suramin and histones (106 ± 2 nM; n=6) and histones (127 ± 2 nM; n=6) samples. (D) Summary data for area under the curve (AUC; nM thrombin) in control (143732 ± 2498 nM; n=6), suramin (139915 ± 2355 nM; n=6), suramin and histones (147896 ± 02968 nM; n=6) and histone (180248 ± 3977 nM; n=6) samples. (E) Summary data for time to peak (min) in control (31 ± 0.8 min; n=6), suramin (29 ± 0.8 min; n=6), suramin and histones (28 ± 0.6 min; n=6) and histone samples (10 ± 0.3 min; n=6). (F) Summary data for velocity (nM thrombin/min) in control (3.0 ± 0.1 nM thrombin/min; n=6), suramin (3.2 ± 0.1 nM thrombin/min; n=6), suramin and histones (3.8 ± 0.04 nM thrombin/min; n=6) and histone (13 ± 0.3 nM thrombin/min; n=6) samples. Data are expressed as mean ± SEM. Ordinary one-way ANOVA with Bonferroni’s correction for multiple comparisons; *P<0.05*.

### Suramin prevents disruption of endothelial-dependent vasodilation and endothelial cell calcium overload caused by histones

Endothelial-dependent vasodilation of small arteries in response to nitric oxide and other hyperpolarizing stimuli is essential for the regulation of regional blood flow to meet metabolic demands. Disruption of endothelial-dependent vasodilation is considered the hallmark of endothelial dysfunction. We previously demonstrated that histones induce aberrant endothelial cell calcium responses that disrupt normal vasodilatory signals in small mesenteric arteries (10, 47). Here, we used video-edge detection to record the vasodilatory responses of pressurized, resistance-sized arteries from the mouse mesenteric circulation to the exposure of the endothelial-dependent vasodilator NS309 (0.1 μM to 1 μM) before and after intraluminally perfusing histones (10 μg/mL) through the vessel in the presence and absence of suramin (50 μM) (Fig. 3 *A* and *B*). Vasodilation to 1 μM NS309 after 30 minutes of histone exposure was reduced to 33 % of the pre-histone control dilation. Vasodilatory function was completely preserved during this same experiment while in the presence of suramin (50 μM). Because lung injury is a significant concern in conditions characterized by high levels of histones such as ARDS (19), we also studied vascular preparations from small mouse pulmonary arteries. These blood vessels were surgically opened on one side to expose the endothelial cell layer for direct measurement of a fluorescent calcium indicator using confocal microscopy (Fig. 3 *C* and *D*). Similar to our prior findings in human and mouse mesenteric arteries (20), we found that histones (10 μg/mL) significantly increased the number of detectable calcium events compared to baseline. The presence of suramin (50 μM) during histone application significantly decreased the number of calcium events; however, the activity was still elevated when compared to the baseline control (Fig. 3 *D*). Together, pre-treatment with suramin completely prevents histone-induced endothelial vasodilatory dysfunction in pressurized arteries and significantly decreases aberrant calcium signaling caused by exposure to histones.

**Figure 3.**
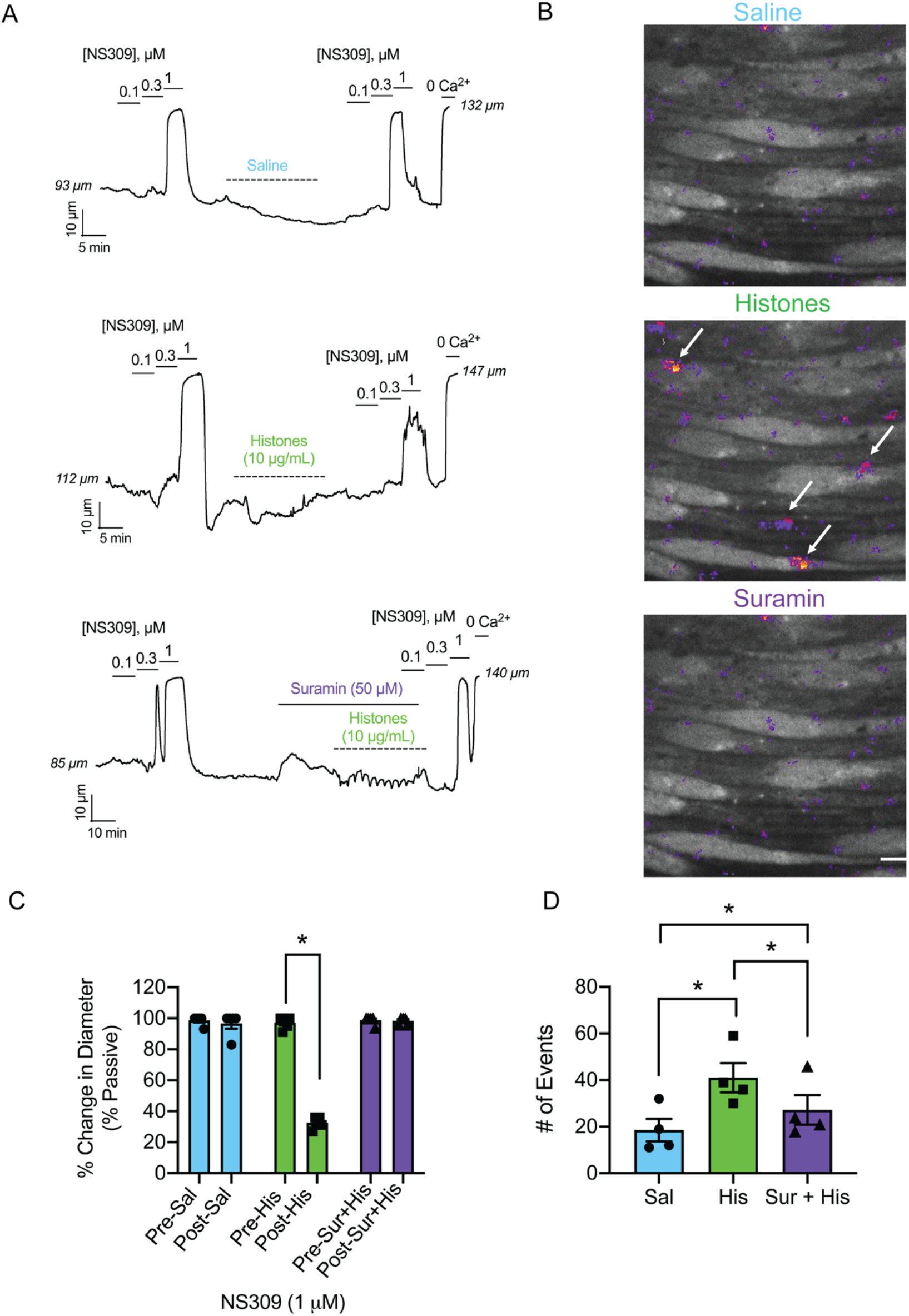
Suramin prevents endothelial dysfunction and calcium overload caused by histones. (A) Representative tracings of pressurized (80 mm Hg), third-order, mouse mesenteric arteries. Histones (10 μg/mL) or saline (control) was flowed through the lumen at 2 μL/min (<5 dynes/cm^2^) for 30 minutes. Dilations to the endothelial-dependent vasodilator NS309 (0.1; 0.3; 1 μM) pre-flow and post-flow were recorded. In one subset of experiments suramin (50 μM) was superfused abluminally for 10 minutes prior to and then continuously during histones flow. Maximal dilations were elicited at the end of the experiments using 0-Ca^2+^ PSS. (B) Paired summary data of percent dilation to 1 μM NS309 pre-flow and post-flow of saline (pre-sal 99 ± 1 vs. post-sal 97 ± 3 %; n=5; n.s.), histones (10 μg/mL) (pre-his 97 ± 2 vs. post-his 33 ± 2 %; n=5; * *P<0*.05; Paired Student’s T-test), and suramin (50 μM) with histones (pre-sur+his 99 ± 1 vs. post-sur-his 98 ± 1 %; n=5; n.s.). (C) Representative images from *en face* mouse pulmonary arteries loaded with Fluo-4 (10 μM) on a spinning disk confocal microscope. All images are from the same field of view recorded over 2 minutes. Arrows indicate large histone-induced calcium event F/F0 ROIs. (D) Summary data of the paired total number of events per field after saline (control; 19 ± 5 events; n=4), histones (His; 10 μg/mL; 41 ± 6 events; n=4), and suramin (50 μM) and histones (Sur+His; 27 ± 6 events; n=4) application. Significant differences were determined using a repeated measures one-way ANOVA test with a Holm-Sidak correction for multiple comparisons for all three groups; *P<0.05*. Data are represented as mean ± SEM. Scale bar = 10 μm.

### Suramin prevents adhesion molecule expression, neutrophil recruitment, and pulmonary endothelial barrier disruption caused by histones

The results of our biochemical and *in vitro* work provided a rationale for an *in vivo* model of histone toxicity. Circulating histones can injure platelets, erythrocytes and vascular endothelial cells from multiple tissue beds. However, in humans and animal models of trauma, lung tissue is particularly vulnerable to circulating damage associated molecular pattern proteins (19). Therefore, we next tested the hypothesis that suramin would prevent the increase in circulating biomarkers, endothelial cell activation, and influx of inflammatory cells into the lungs caused by histone infusion (45 mg/Kg). To specifically assess the endothelial effects of suramin after histone exposure, we freshly isolated mouse pulmonary endothelial cells and measured their adhesion molecules using flow cytometry. The extent of neutrophil migration into the lungs after histone exposure was also quantified. After 24 hours, lung tissue was dissociated, and the frequency of neutrophils determined by expression of CD11b and Ly6G by flow cytometry. Treatment with histones resulted in a statistically significant increase in the frequency of neutrophils in the lung (Fig. 4 *A*). Lung endothelial cell Intracellular Adhesion Molecule-1 (ICAM-1) expression was also significantly increased by histones (Fig. 4 *B*). Suramin ameliorated these histone-induced effects, causing a significant reduction in the frequency of neutrophils and expression of endothelial cell ICAM-1. Lastly, compared to controls, treatment with suramin before histone exposure significantly reduced the levels of circulating adhesion proteins shed by activated endothelial cells (E-selectin, P-selectin, ICAM-1, and PECAM-1; Fig. 4 *C*).

**Figure 4.**
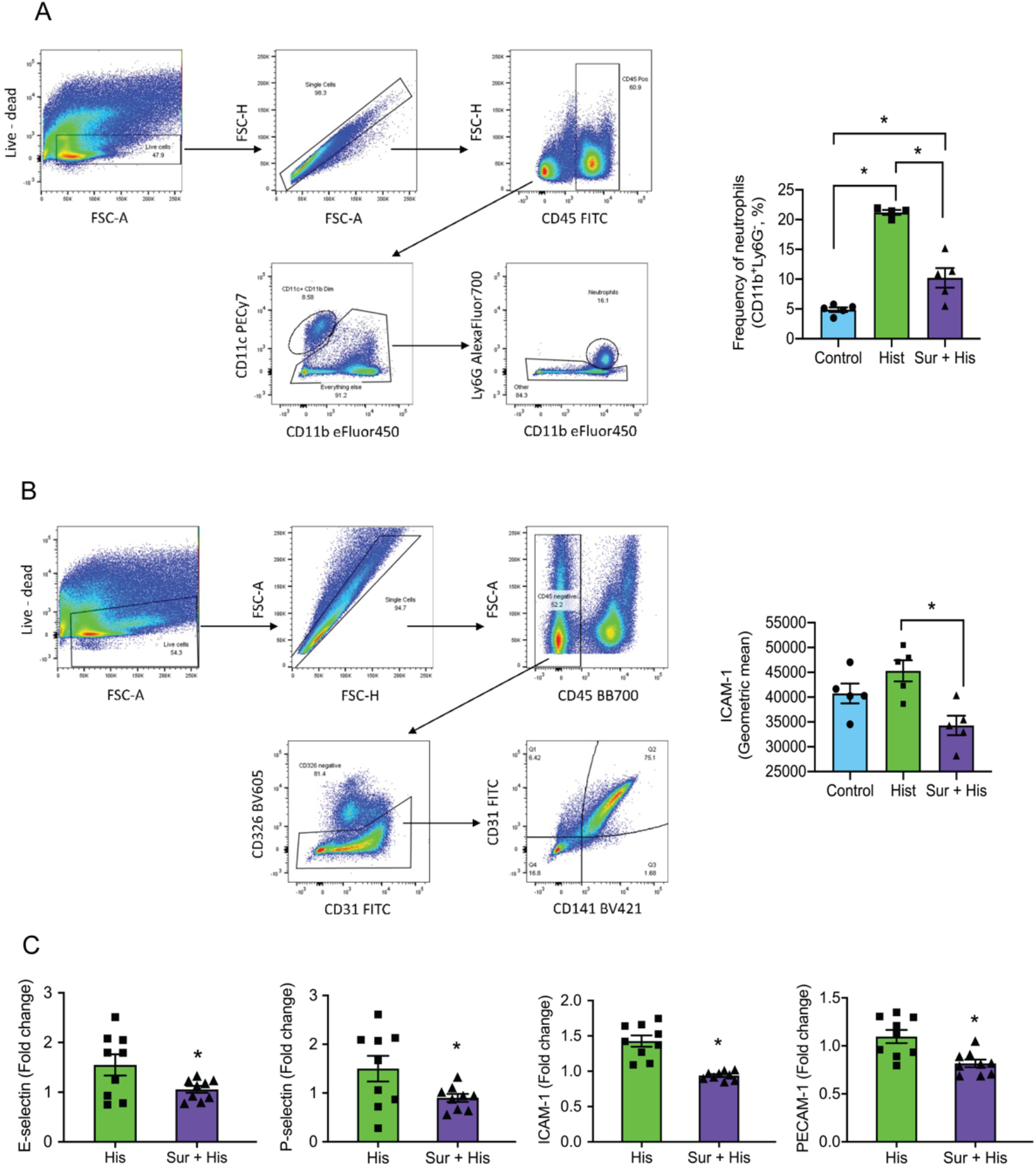
Histone-induced neutrophil recruitment and adhesion molecule expression is blocked by suramin. (A) Live cells were gated and doublets excluded (FSC-A vs FSC-H). CD45^+^ cells were selected and CD11c^−^ cells identified. Neutrophils (CD11b^+^Ly6G^+^) were identified, and the frequency of neutrophils per live cells determined. Summary data of the frequency of neutrophil levels in lung tissue 4 hours after saline (Control; 300 μL; 4.8 ± 0.4 frequency; n=5), histones (Hist; 45 mg/Kg; 21 ± 0.4 frequency; n=4), or suramin (50 mg/Kg) and histone injection (Sur+His; 10 ± 1.6 frequency; n=5). (B) Live cells were gated and doublets excluded (FSC-H vs FSC-A). CD45^−^ cells were selected and CD31^+^CD326^−^ cells identified. Endothelial cells (CD31^+^CD141^+^, Q2) were assessed for CD54 expression (geometric mean intensity). Summary data for endothelial ICAM-1 (CD54) expression geometric means (GM) in lung tissue 4 hours after saline (Control; 40743 ± 1999 GM; n=5), histones (Hist; 45 mg/Kg; 45314 ± 2126 GM; n=5), or suramin (50 mg/Kg) and histone injection (Sur+His; 34288 ± 1957 GM; n=5). Data are expressed as mean ± SEM. Two-way ANOVA with Bonferroni’s correction for multiple comparisons; *P<0.05*. (C) Suramin decreases histone-induced adhesion molecule levels in serum. Adhesion molecule levels measured in serum with a multiplex immunoassay (Luminex). Samples were collected 4 hours after histones (His; 45 mg/Kg) and histone with suramin (50 mg/Kg) injection (Sur+His). Suramin reduced circulating levels of (A) E-selectin, P-selectin, ICAM-1 and PECAM-1. Data are expressed as fold change, mean ± SEM. Student’s T-test; *P<0.05*.

### Suramin prevents lung injury and improves survival after exposure to histones

To assess clinically relevant outcomes, we next tested whether suramin would prevent death and lung injury *in vivo* after histone exposure. We randomized mice to one of 4 experimental groups: saline; saline and histones (75 mg/Kg); suramin (20 mg/Kg) and histones; or suramin (50 mg/Kg) and histones (Fig. 5 *A*). Survival was monitored and updated every minute for the 35-minute duration of the study. Groups were compared using a Mantel-Cox analysis and Mantel-Haenszel for the Hazard Ratio. Seventy percent (70 %) of animals receiving 75 mg/Kg of histones alone died abruptly within 10 minutes and showed symptoms such as bleeding from the nose, pink frothy sputum and signs of respiratory distress, and only 20 % survived the 35-minute period. In contrast, 100 % of animals receiving suramin at the higher dose of 50 mg/Kg with the lethal dose of histones survived when compared to the histone group (1.959 to 35.09 95 % CI; 8.29 Hazard Ratio; *P<0.05; \**). The lower dose of suramin (20 mg/Kg) also provided a modest survival benefit (36 %) when compared to the high dose (1.359 to 28.71 95 % CI; 6.25 Hazard Ratio; *P<0.05; *)* but not to the histone only group (0.9702 to 9.410 95 % CI; 2.49 Hazard Ratio; n.s.). In a separate set of mice exposed to histone infusion (45 mg/Kg) in the presence or absence of the higher dose of suramin (50 mg/Kg), we found that suramin reversed intra-alveolar hemorrhage visible on histology, and elevation in cell counts and protein measured in broncho-alveolar lavage fluid (Fig. 5 *B* *and C)*. Pulmonary barrier breakdown induced by histones, quantified as the extravasation of 70-kDa FITC-labeled dextran, was significantly decreased by suramin. Suramin also blocked extravasation of the labeled dextran in renal tissue caused by histones (Fig. 5 *D*).

**Figure 5.**
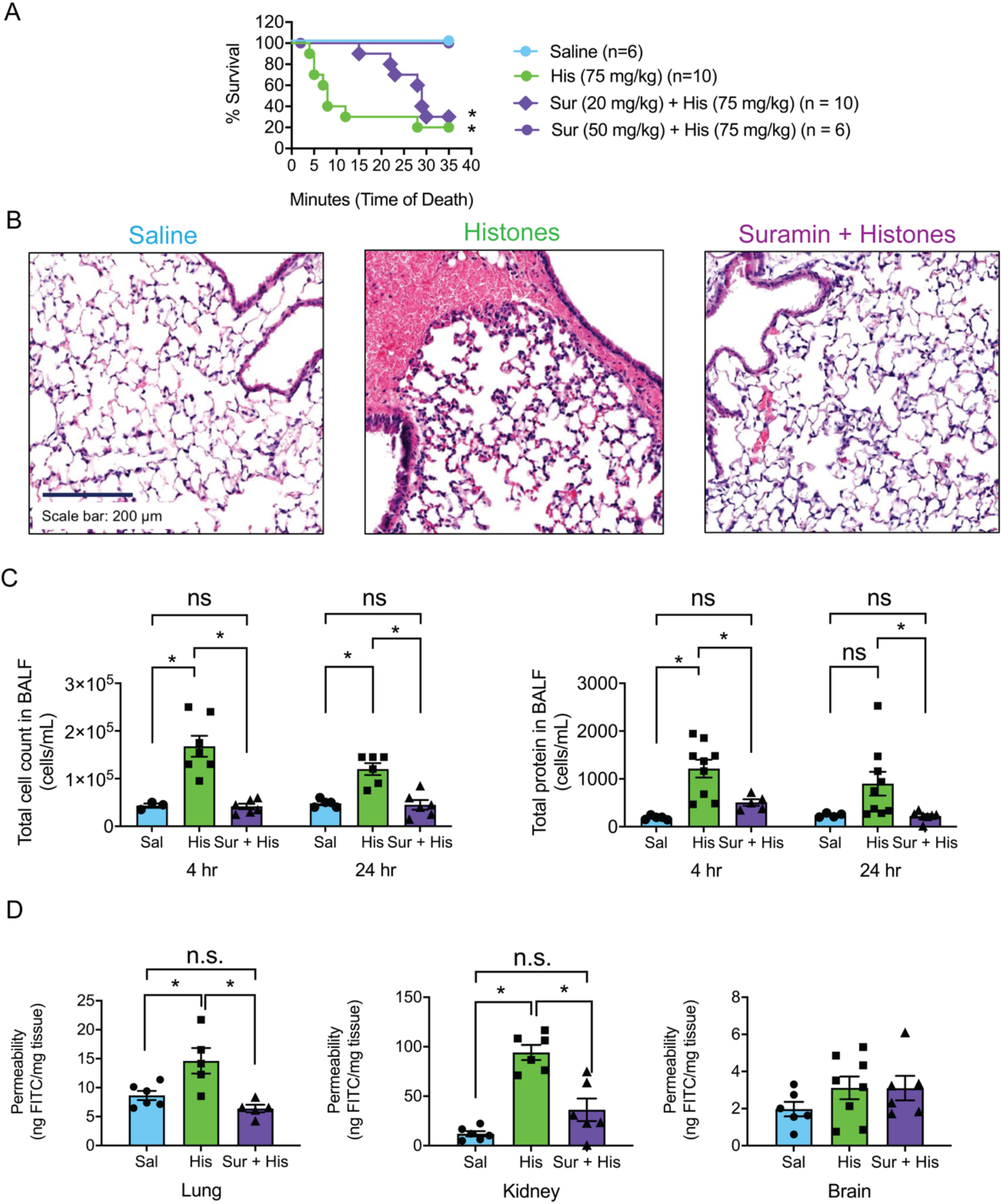
Suramin improves survival and prevents lung injury and edema caused by histones. (A) Saline injected (control; n=6), a lethal dose of histones (His; 75 mg/Kg; n=10), and a lethal dose of histones with suramin (Sur+His; 20 mg/Kg n=11 or 50 mg/Kg n=6) was injected into mice and survival was recorded over the course of 35 minutes. Suramin was injected intraperitoneally. Mantel-Cox test; *P<0.05*; *, for each group compared to saline. (B) Representative images of hematoxylin and eosin stain (H&E) of a histological section of paraffin-embedded fixed lung tissue from a mouse from each group. The dark blue color denotes cell nuclei, light pink extracellular matrix, and the red erythrocytes. Scale bar = 200 μm. (C) Summary data of the total, non-differentiated cell counts in the bronchial-alveolar lavage fluid (BALF) at 4 hours after saline (control; 43333 ± 4410 cells/mL; n=3), histones (His; 45 mg/Kg; 167847 ± 22008 cells/mL; n=7), or suramin (50 mg/Kg and histone injection (Sur+His; 41667 ± 5725 cells/mL; n=6) and 24 hours after saline (control; 48000 ± 3742 cells/mL; n=5), histones (His; 45 mg/Kg; 120000 ± 12649 cells/mL; n=6), or suramin (50 mg/Kg) and histone injection (Sur+His; 45000 ± 10247 cells/mL; n=6). Summary data for the total protein leakage into the BALF at 4 hours after saline (control; 188 ± 19 μg/mL; n=5), histones (His; 45 mg/Kg; 1215 ± 186 μg/mL; n=9), or suramin (50 mg/Kg) and histone injection (Sur+His; 507 ± 66 μg/mL; n=5) and 24 hours after saline (control; 239 ± 21 μg/mL; n=5), histones (His; 45 mg/Kg; 901 ± 249 μg/mL; n=9), or suramin (50 mg/Kg) and histone injection (Sur+His; 225 ± 39 μg/mL; n=7). Data are expressed as mean ± SEM. Two-way ANOVA with Bonferroni’s correction for multiple comparisons; *P<0.05*. (D) Summary data for FITC-dextran extravasation using the modified Mile’s Assay in lung, kidney, and brain tissue from saline (control), histones (45 mg/Kg) and suramin (50 mg/Kg) and histone (Sur+His) injected mice at 4 hr. Lung permeability in saline (control; 8.7 ± 0.8 ng FITC/mg tissue; n=6), histones (His; 14.6 ± 2.2 ng FITC/mg tissue; n=5), and suramin and histone (Sur+His; 6.4 ± 0.7 ng FITC/mg tissue; n=5) injected mice. Kidney permeability in saline (control; 12 ± 2.7 ng FITC/mg tissue; n=6), histones (His; 94 ± 7.7 ng FITC/mg tissue; n=6), and suramin and histone (Sur+His; 36 ± 11 ng FITC/mg tissue; n=6) injected mice. Brain permeability in saline (control; 1.9 ± 0.4 ng FITC/mg tissue; n=6), histones (His; 3.1 ± 0.6 ng FITC/mg tissue; n=6), and suramin and histone (Sur+His; 3.1 ± 0.6 ng FITC/mg tissue; n=6) injected mice. Data are expressed as mean ± SEM. Two-way ANOVA with Bonferroni’s correction for multiple comparisons; *P<0.05*.

**Figure 6.**
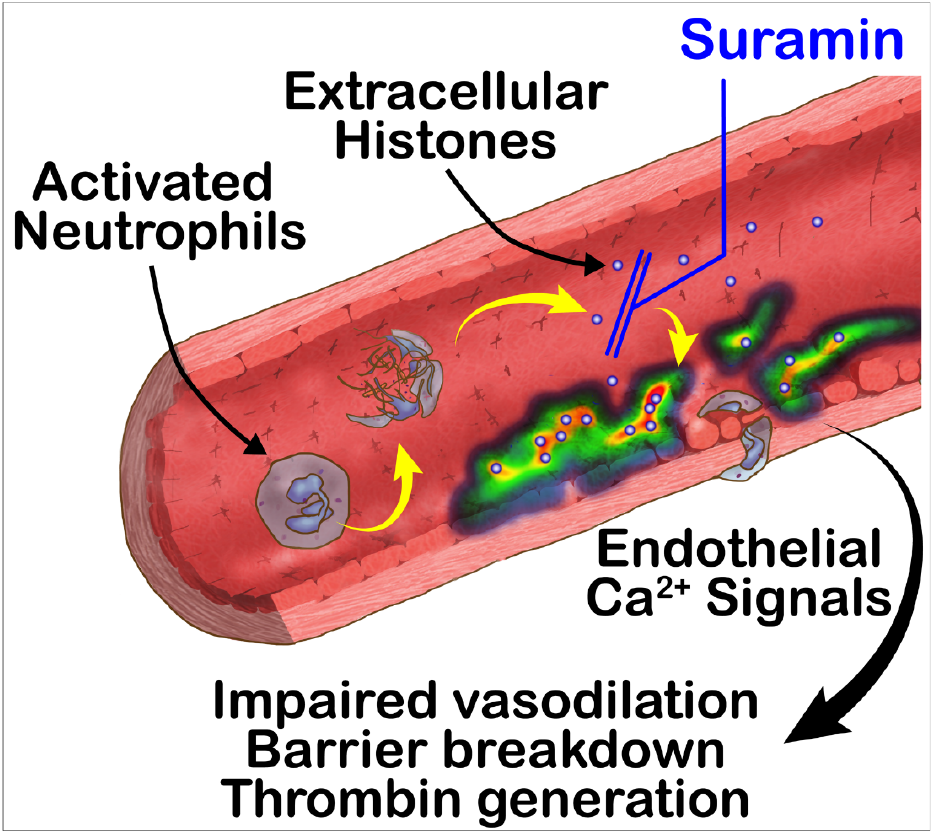
Schematic overview of the inhibitory effects of suramin on histones-endothelial interactions. Suramin prevents histone-induced calcium overload and death, impaired vasodilation, endothelial barrier breakdown, neutrophil migration, adhesion molecule expression, and thrombin generation in vascular endothelial cells.

## Discussion

Histones can activate and damage vascular cells through several mechanisms that are not fully understood. Histones can activate ion channels, observed by membrane potential and current recordings in endothelial cells and other cells (19, 54, 55). With prolonged exposure (minutes to hours), or at high concentrations, histones – and histone H4 in particular – can also damage lipid bilayers in any cell type, including endothelial cells, and act as cell-penetrating proteins (11, 22, 31). Histones can also engage innate immune responses leading to prothrombotic activation of endothelial cells (27, 46, 56) or pyroptosis (6, 24–26). All these mechanisms can contribute to acute endotheliopathy. Here, we demonstrate that not only does suramin form a stable complex with histone proteins, but also, this neutralizing effect completely prevents histone-induced endothelial dysfunction and mortality. Suramin has been used for over 100 years as an anti-parasite and anti-cancer agent, and, importantly, is considered among the safest and most effective drugs for health care by the World health organization. This discovery of a new mechanism of action for a widely available and easily administered drug, as a blocker of deleterious histone effects, provides a tantalizing target for the potential clinical therapeutic use of suramin in acute immuno-vascular and thrombo-inflammatory conditions.

Our results provide new insight into the pathophysiological outcomes of histone-induced organ injury. We provide the first demonstration that in native, pulmonary artery preparations, histones elicit calcium-mediated events similar to those we previously observed in mesenteric resistance arteries from human and mouse. We also found that histone infusion caused endothelial barrier breakdown of small blood vessels in both kidney and lung, with increased extravasation of the 70-kDa dextran, but not brain. This is consistent with other evidence that pulmonary and renal (6, 17, 19) tissue beds are highly sensitive to histone-induced injury. It was recently shown that histones increased paracellular permeability in the hippocampus but not cortical brain regions (23). It is possible that we missed these regional cerebrovascular effects because we quantified vascular leak for the entire brain and not specific regions, or because we used a 70-kDa tracer rather than a smaller sized dextran which would more specifically target blood-brain barrier permeability. Here, our focus was on lung injury, with results in freshly harvested lung cells, isolated vascular preparations, and *in vivo* models uniformly supporting model in which histones activate endothelial cells to increase cellular adhesion molecule expression, in conjunction with increased release of circulating adhesion molecules. These changes in the pulmonary vascular endothelium result in increased neutrophil recruitment to the lungs.

Importantly, we also demonstrate a new, endothelial-dependent mechanism by which histones increase thrombosis. Prior studies have shown that histones can increase plasma thrombin generation in purified systems by reducing thrombomodulin-dependent protein C activation (57). Here, we provide new evidence that histones can rapidly activate endothelial cells directly to promote thrombin generation. We show that on endothelial cells, in the absence of added tissue factor (TF) or phospholipids, thrombin formation occurs slowly, but when histones are applied, thrombin generation is accelerated, with a lag time of <5 minutes. This reaction was blocked by suramin. The time course of this reaction, occurring minutes after histone exposure, is not consistent with known pro-coagulant responses of endothelial cells to histones, such as release of VWF (56), upregulation of TF (27), or downregulation of thrombomodulin mRNA and surface antigens (46) which occur 1 to 8 hours after exposure. Thus, rapid phosphatidylserine translocation, possibly due to TMEM16f activation (58, 59), coupled with mobilization of “cryptic” TF in the endothelial cell membrane, likely drive the rapid reactions we observe. Understanding the effectors of rapid procoagulant responses to histones may improve targeted therapies to protect against excessive thrombosis in inflammatory conditions.

Suramin offers several advantages over other therapeutic strategies to prevent histone-medicated vascular injury. Polyanions such as heparin can neutralize histones and prevent histone-mediated cytotoxicity (42, 60–62). Heparin improves outcomes in some patients with sepsis (60) or COVID-19 (63, 64), but the mechanisms are not fully understood. Furthermore, heparin cannot be safely used in all patients, such as those requiring surgical procedures, because of the risk of hemorrhagic bleeding. Unlike heparin which may require continuous infusion, suramin dosing is infrequent (once per week), and it is inexpensive, well-tolerated, and extensive experience with this drug has shown that it has an excellent safety profile. It is readily available world-wide, at a low cost. Other naturally-occurring substances, such as pentraxin 3 (65), activated protein C (APC) (35), C1 esterase inhibitor (66), and inter-α-inhibitor proteins (39); anti-histone antibodies (3, 4, 42) and synthetic polyanions (22, 67), can also neutralize excessive histones to prevent toxicity, but these drugs are either not approved or not available for human use. Albumin or fresh frozen plasma may have benefits in trauma and sepsis, in part due to histone binding (68), but blood products are limited resources with high costs compared to suramin.

Taken together, these results provide evidence supporting the use of suramin in trauma and sepsis. This particularly important in the context of the unprecedented global public health crisis caused by the novel SARS-CoV-2 virus, because histone levels are elevated in individuals with COVID-19 (18, 69), and endotheliopathy and thromboinflammation secondary to NETs drives progression from systemic inflammation to organ failure and death (1, 2). Of note, suramin may have other mechanisms of therapeutic action in viral illness. Polyanion inhibitors have been used to block viruses that require cell surface sugars such as heparan sulfate (HS) to infect humans, which include HIV, Ebola, Zika, and SARS. These inhibitors include naturally occurring therapeutic polyanions such as heparin, synthetic polyanions, such as suramin, or modified cyclodextrins (70). Heparin is the subject of over a dozen registered clinical trials for SARS-CoV-2 (clinicaltrials.gov) and has efficacy in COVID-19 (64). Most of these trials are testing injectable unfractionated heparin or low molecular weight heparin at prophylactic or therapeutic anticoagulant doses. Other trials are using intranasal or nebulized heparin in an attempt to block viral entry, because heparin can serve as a decoy for the target cell heparan sulfate needed for optimal interaction between viral spike protein and ACE2. Heparin is also believed to have immunomodulatory and endothelial protective effects, based on evidence of prior benefit in sepsis from other causes, and histone binding may be an important therapeutic mechanism of action for heparin (60, 71). Suramin is a competitive inhibitor of heparin, and it has been suggested – but not established – that suramin shares a mechanism of action against SARS-CoV-2 by acting as a decoy for heparan sulfate that can block spike protein and ACE2 interactions (72). Suramin also inhibits the main protease needed for SARS-CoV-2 infection (73), and it is the subject of at least one COVID-19 clinical trial at the time of this submission (clinicaltrials.gov).

In summary, we demonstrate a new and previously unreported mechanism of action for an old drug: suramin blocks cytotoxic effects of histones and prevents histone-induced vasodilatory dysfunction, endothelial cell activation, thrombin generation, lung injury, and mortality in mice. Thus, our results provide a mechanistic basis and rationale for clinical trials of suramin as a repurposed treatment that can be rapidly deployed to prevent endothelial injury and excessive blood clotting in conditions associated with high circulating histone levels such as trauma and sepsis.

## Materials and Methods

### Animals

Male C57BL/6 J mice (12 weeks old; ~30 g) were purchased from The Jackson Laboratory (Bar Harbor, ME). All animals were maintained on 12-hour light/dark cycle and standard diet and water *ad libitum* in an AALAC-accredited facility. All animal experiments were approved by the University of Vermont’s Institutional Animal Care and Use Committee, in accordance with the recommendations in the Guide for the Care and Use of Laboratory Animals of the National Institutes of Health, and efforts were made to minimize suffering.

### *In vivo* histone studies

Mice were anesthetized with 2.5% isoflurane, and calf thymus histones (45 mg/Kg in sterile saline; Roche® distributed by Sigma-Millipore; #10223565001; containing H1, H2a, H3, H4 histones) or sterile saline was injected via the retroorbital sinus. Prior to histone injection, suramin (Adipogen; #AG-CR1-3575) was injected intraperitoneally (20 mg/Kg; 50 mg/Kg). The approved dose of suramin is 1 g for adults and 10–15 mg/Kg for children (http://home.intekom.com/pharm/bayer/suramin.html). We used 20 mg/Kg as the highest human reference dose. Mice were divided into the following treatment groups: control (sterile saline; i.v.), histones (45 mg/Kg; i.v.), histones + suramin (20 mg/Kg; 50 mg/Kg; i.p.) (administered 1 hour prior to histone infusion). At 24 hours after treatment, mice were terminally anesthetized and euthanized. Additional survival studies were also performed in mice that received a lethal dose of histones (75 mg/Kg; i.v.) (1); the survival rates were determined every five minutes, for 1 hour.

To assess for markers of activated endothelium, blood was collected via cardiac puncture in BD Microtainer® blood collection tubes (BD Biosciences). Sera was obtained by centrifugation (1,300 x g, 10 min) and frozen. Thawed sera were diluted two-fold and cardiovascular markers were measured using a Milliplex mouse cardiovascular disease magnetic bead panel (Millipore-Sigma; #MCVD1MAG-77K). Data were acquired using the Bio-Plex suspension array system and Bio-Plex Manager software.

To assess inflammation in the lung, bronchoalveolar lavage (BAL) fluid was collected and analyzed for total number of leukocytes and total protein. Euthanized mice were tracheotomized with an 18-gauge cannula and lavaged with 1 mL Dulbecco’s phosphate-buffered saline (Life Technologies, Carlsbad, CA). Lavage fluid was centrifuged (1,300 x g, 10 min) and cell-free supernatants were snap-frozen for total protein analysis using the Pierce™ BCA protein assay kit (Thermo Scientific; #23227). The pellet was resuspended with 400 μL of PBS and total leukocyte count was measured via a hemocytometer (Neubauer chamber). Intact lungs were also assessed for histological changes. Lungs were inflation-fixed at 20 cm H_2_O pressure with buffered formalin for 24 hours, embedded in paraffin, sectioned, and stained with hematoxylin and eosin.

### Calcium imaging on pulmonary arteries

Calcium imaging in the native endothelium of mouse pulmonary arteries was performed as previously described (74). Briefly, 4^th^-order (~50 μm) pulmonary arteries were pinned down *en face* on a Sylgard block and loaded with Fluo-4-AM (10 μM) in the presence of pluronic acid (0.04 %) at 30° C for 30 minutes. Fluo-4 was excited at 488 nm with a solid-state laser and emitted fluorescence was captured using a 525/36-nm band-pass filter. Images were acquired at 30 frames per second with Andor Revolution WD (with Borealis) spinning-disk confocal imaging system (Oxford Instruments, Abingdon, UK) comprised of an upright Nikon microscope with a 60X water dipping objective (numerical aperture 1.0) and an electron multiplying charge coupled device camera (iXon 888, Oxford Instruments, Abingdon, UK). Ca2+ signals were analyzed using the custom-designed SparkAn software (47, 74). A region of interest defined by a 1.7-μm^2^ (5×5 pixels) box was placed at a point corresponding to peak event amplitude to generate a fractional fluorescence (F/F_0_) trace. F/F_0_ traces were filtered using a Gaussian filter and a cutoff corner frequency of 4 Hz. The number of Ca^2+^ events was auto-detected using a detection threshold of 0.3 F/F_0_ in SparkAn (custom software, Dr. Adrian Bonev, Burlington, VT). Each data point indicates one field of view from one pulmonary artery.

### *In vivo* assessment of pulmonary vascular permeability

Vascular permeability to solutes was assessed by measuring extravasation of FITC-labelled 70-kDa dextran. Mice were given a retroorbital injection of 70-kDa FITC-dextran (100 μL of 3 mg/mL) and then after 30 minutes transcardially perfused with PBS to eliminate any remaining FITC-dextran in circulation. Lungs were isolated and homogenized in 1 mL RIPA buffer and centrifuged at 12,000 x g for 20 minutes. The concentration of 70-kDa FITC-dextran in the supernatant was detected via fluorescence measurement (excitation 490 nm, emission 520) and interpolation from a standard curve of known concentrations of FITC-dextran. Results are presented as ng of FITC-dextran per mg of protein in the supernatant.

### Fluorescence spectroscopy

Fluorescence spectroscopy was used to determine the equilibrium dissociation constant (K_d_) values for the histone-suramin complex. The changes in intrinsic suramin fluorescence emission were measured with a microplate reader (BioTek; Winooski, VT) at 25 °C. The samples were excited at 315 nm and the emission spectrum was measured between 370-480 nm. Histones did not show spectral overlap in that range (Supplemental Fig. 1 *A*). The titration was performed stepwise with a suramin stock concentration (1 μM) in assay buffer containing 50 mM HEPES, 100 mM NaCl and 2 mM CaCl_2_ (pH 7.4); fluorescent measurements were performed after each titration with histones (0 to 8 μM). After normalization of the fluorescence emission signal, Kd for each suramin-histones complex was estimated by nonlinear curve fitting with a sigmoidal dose– response function using GraphPad 7 software (GraphPad, San Diego, CA). The percentage of bound suramin-histones was plotted against the concentration of free histones.

### Molecular modeling

#### Model Preparation

All the models were constructed using the Desmond/Maestro program (v2016-3 Schrödinger, Inc.) using the System Builder in Maestro. Each model contained a complete histone octamer (PDB: 5XF3) with or without DNA and six suramin molecules that were arbitrarily placed at a minimum distance of 15 Å from the proteins. The SPC water model was employed to solvate the complexes, with counter ions and 0.12 M NaCl, 0.047 M KCl, 0.025 M CaCl_2_, and 0.012 M MgCl_2_. The construct with a DNA-bound histone has a total of 197,122 atoms in a periodic box of ~123×128×127 Å^3^, while the one with a DNA-free histone has 180,860 atoms in a box of ~ 120×122×124 Å^3^.

#### Simulation setup

All simulations were performed in the Desmond program with the OPLS3 force field in the NPT ensemble (1.01325 bar, 310 K, Martyna-Tobias-Klein coupling scheme) with a time step of 2 fs(75, 76). The particle mesh Ewald technique was used for the electrostatic calculations. The Van der Waals and short-range electrostatics were cut off at 9.0 Å. Hydrogen atoms were constrained using the SHAKE algorithm. Each simulation has two 700-ns replicas.

#### Visualization and Analysis

PyMOL (v2.5, Schrödinger, Inc.) and Visual Molecular Dynamics (VMD, http://www.ks.uiuc.edu/Research/vmd/) were used for the structure visualization of the simulations, while the simulation analysis panel was carried out in Maestro.

### Fresh isolation of lung cell populations and flow cytometry

Mice were euthanized using sodium pentobarbital. Lungs were inflated with an enzymatic digestion buffer (Dubecco’s modified Eagle’s medium [DMEM], 1 mg/mL collagenase type IV [Invitrogen], and 0.2 mg/mL DNase I [Sigma]), after which they were dissected away from the trachea and heart and incubated in 5 mL enzymatic digestion buffer in a 50 mL conical tube for 30 min at 37°C under agitation at 200 rpm. After the 30-min incubation, 25 mL of PBS was added and the samples were vortexed for 30 seconds. The resulting cell suspension was passed through a 70-μm filter and washed in PBS. Red blood cells were lysed using Gey’s solution and washed in PBS–2% fetal calf serum, after which cells were counted and resuspended for flow cytometry experiments.

For flow cytometry, nonspecific antibody binding was blocked by incubating 1×10^6^ cells with FcBlock anti-CD16/32 (BD Biosciences; #553141). After washing, cells were stained at 4°C in PBS–2% FCS containing 0.1% sodium azide. Reagents and antibodies used in these experiments were as follows: Live/Dead™ (Invitrogen; #L23105), CD45-FITC (eBioscience; #MCD4501), CD11c-PECy7 (BD Biosciences; #561022), CD11b-eFluor450 (eBioscience; #48-0112-82), Ly6G-Alexa Fluor700 (BD Biosciences; #561236), CD45 BB700 (BD Biosciences; #566440), CD326 BV605 (BD Biosciences; #740389), CD31-FITC (BD Biosciences; #558738), CD141 BV421 (BD Biosciences; #747647). Data were collected on a BD LSRII flow cytometer (BD Biosciences) and analyzed using FlowJo (TreeStar, Ashland, OR).

### Calibrated automated thrombinography

Human endothelial cells (EA.hy926; ATCC® CRL-2922™) were incubated with histones (50 μg/mL), suramin (50 μM), histones + suramin, or Dulbecco’s Modified Eagle Medium alone for 4 hr at 37°C, 5% CO_2_. The cells were released from the tissue culture wells with trypsin and subjected to centrifugation (170 x g, 7 min). Cell pellets were washed one time by resuspension in 20 mM HEPES, 0.15 M NaCl (pH 7.4) (HBS) followed by centrifugation. The final cell pellets were resuspended in HBS and adjusted to a final concentration of 1×10^7^/mL.

Thrombin generation was assessed using a modified calibrated automated thrombogram. Plasma was thawed at 37°C in the presence of corn trypsin inhibitor (0.1 mg/mL final concentration), and incubated with the thrombin substrate Z-Gly-Gly-Arg 7-amido-4-methylcoumarin hydrochloride (0.42 mM) (Bachem AG, Switzerland) and CaCl_2_ (15 mM) (3 min, 37°C). The reactions were initiated by the addition of relipidated tissue factor_1-242_ (6.5 pM) (a gift from Dr. R. Lunblad, Baxter Healthcare Corp.) and synthetic vesicles consisting of 80% phosphatidylcholine and 20% phosphatidylserine (PCPS) (20 μM), or EA.hy926 cells (2×10^5^). Fluorescence was measured (ex = 370 nm/em = 460 nm) for 1 hour with a Cytation 3 imaging reader (BioTek, Winooski, VT). Changes in fluorescence were converted to thrombin concentrations using a calibration curve created from sequential dilutions of human thrombin. If no change in fluorescence was noted after 60 min, the lag time for the sample was defined as >60 min.

### Statistics

Data sets were first tested for normal distribution using the Kolmogorov-Smirnov method to determine the appropriate parametric or non-parametric test with which to proceed. Data were analyzed by two-tailed unpaired or paired Student’s T-test, Mann-Whitney U-test, Wilcoxon rank text, one-way or two-way ANOVA and Bonferroni post-hoc test, or the Mantel-Cox test for survival, using GraphPad Prism 7.04 (GraphPad Software, Inc, La Jolla, CA). A P value less than 0.05 was considered statistically significant.

## Data sharing

All data, documentation and code generated or utilized for this paper is available by request.

## Acknowledgments

We thank Dr. Roxana Del Rio Guerra and the University of Vermont Larner College of Medicine Flow Cytometry and Cell Sorting Facility for extensive support on all flow cytometric experiments as well for instrumentation. Imaging work was performed at the Microscopy Imaging Center at the University of Vermont (RRID# SCR_018821). Research reported in this publication was supported by the Totman Medical Research Trust (to M.T.N.), the European Union Horizon 2020 Research and Innovation Programme (Grant Agreement 666881, SVDs@target, to M.T.N.), as well as grants from the National Institute of Neurological Disorders and Stroke (NINDS) and National Institute of Aging (NIA) (R01-NS-110656 to M.T.N.), the National Institute of General Medical Sciences (NIGMS) (P20-GM-135007 to M.T.N and R35-GM-144099-01 to K.F.), and the National Heart, Lung, and Blood Institute (NHLBI) (R01-HL-146914 to S.S., R01-HL-157407 to S.S., R01-HL-142081 to M.E.P, R01-HL-133920 to M.E.P., R35-HL-140027 to M.T.N., and 1OT2HL156812-01 to K.F.); the the Department of Defense/ The Henry M. Jackson Foundation for the Advancement of Military Medicine (HU001-18-2-0016 to K.F.); the Office of the Director, National Institutes of Health (S10-OD-026843 to J.E.B.); the National Institute of Allergy and Infectious Diseases (R03-AI153902 to J.E.B); and the American Chemical Society (ACS-PRF-58219-DNI to J.L. and Y.T).

---

**Supplemental Figure 1.**
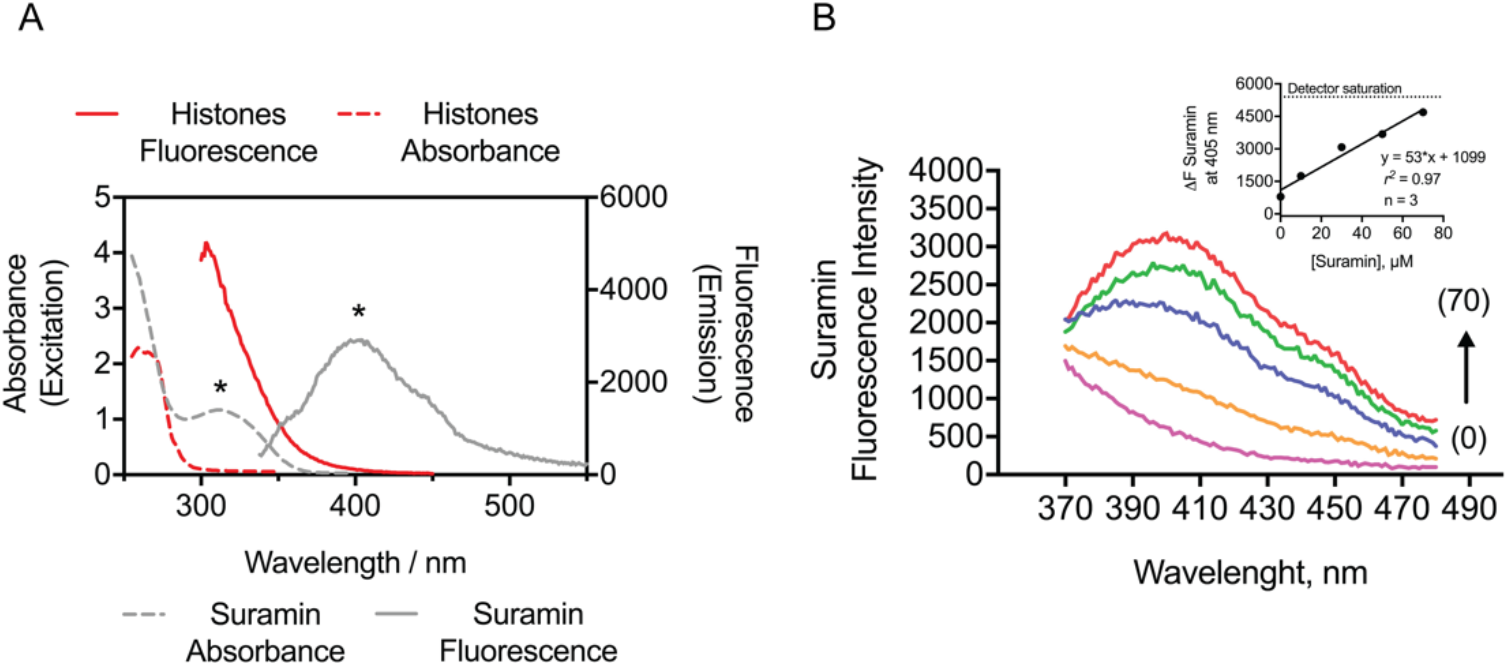
Emission (—) and excitation (- - -) spectra of suramin (*) and histones in HEPES buffer (pH 7.4). λ_exc_ = 315 nm and λ_em_ = 405 nm were used for the emission and excitation scan, respectively, of suramin fluorescence, whereas λ_exc_ = 278 nm and λ_em_ = 305 nm were used for the emission and excitation scan, respectively, of histones. (A) Dual plot of both histones and suramin fluorescence and absorbance. (B) Suramin intrinsic fluorescence intensity in solution from 0 (0) to 70 μM (70).

**Supplemental video 1**. A molecular dynamics simulation shows electrostatic interactions between suramin molecules (stick representation) and the histone octamer (cartoon representation), with hidden water molecules and counter ions for clarity. Six suramin molecules were first arbitrarily placed in the simulation box at the proximity of histone. At the end of the 700-ns simulation, five of six suramin molecules formed stable contacts with the protein, mainly via salt bridges and hydrogen bonding.

